# Century long fertilization reduces stochasticity controlling grassland microbial community succession

**DOI:** 10.1101/638908

**Authors:** Yuting Liang, Daliang Ning, Zhenmei Lu, Na Zhang, Lauren Hale, Liyou Wu, Ian M. Clark, Steve P. McGrath, Jonathan Storkey, Penny R. Hirsch, Bo Sun, Jizhong Zhou

## Abstract

Determining the drivers underlying ecological succession is a fundamental goal of ecological research and essential for predicting ecosystem functioning in response to human-induced environmental changes. Although various studies have examined the impacts of nitrogen (N) addition on plant and microbial community diversity, structure and activities, it remains unknown how long-term anthropogenic fertilization affects the ecological succession of microbial functional guilds and its underlying community assembly mechanisms. Here, using archived soils, we examined more than a century’s succession in soil microbial functional communities (from 1870 to 2008) from the Park Grass Experiment at Rothamsted Experimental Station, the longest running ecological experiment in the world. Long-term fertilization was found to significantly alter soil functional community structure and led to increasingly convergent succession of soil microbial communities. Meta-analysis indicated that microbial temporal turnover (*w*) was highly time scale-dependent, and the *w* value threshold was estimated as 0.0025 with a threshold time point of approximately 160 years. In addition, the importance of stochastic assembly varied greatly in regulating the succession of different microbial guilds. Fertilization had large to medium effects on reducing ecological stochasticity for microbial guilds involved in carbon (C) fixation and degradation, N fixation and mineralization, and denitrification. This century long-term study elucidated the differing influences of assembly mechanisms on soil microbial functional communities involved in C and N cycling, which could not be derived from taxonomic or phylogenetic approaches.

## Introduction

Ecological succession, the change in community composition and structure over time, is arguably the oldest and most intensively studied concept in animal and plant ecology^1^. Determining temporal succession and its underlying mechanisms are fundamental objectives of ecological research and critical for predicting ecosystem functioning in response to anthropogenic activities. Although microbial communities constitute a large portion of the Earth’s biosphere and are important in maintaining various biogeochemical processes that mediate ecosystem functions, the temporal succession and assembly mechanisms of microbial communities are poorly understood. Recently, using high-throughput metagenomic technologies, various temporal dynamic patterns were reported in microbial communities from various habitats, such as species-time relationship (STR)^2–7^ and time decay relationship (TDR)^8–11^. It is generally considered that the temporal turnover of microbial communities is lower than that of larger organisms due to the unique biology of microorganisms, including their massive population sizes, high dispersal rates, rapid asexual reproduction, and resistance to extinction^10^. However, the mechanisms driving such temporal successional patterns remain controversial. For instance, under resource-rich and low-stress environments, microbial succession was found to be governed primarily by stochastic processes (e.g., dispersal and drift)^12–14^. In contrast, under environments with selective pressures, microbial succession was reported to be controlled by deterministic abiotic filtering, including temperature^15,16^, pH^13^, resource type^7,17^, nutrient availability^18,19^, and/or biotic interactions^20,21^.

A grand challenge in contemporary microbial ecology studies is to link the functional succession of microbial communities to environmental change^19,22^. Most studies are based on taxonomic and/or phylogenetic diversity but not on functional diversity. It is believed that functional traits could have more critical impacts on community assembly^23^, and the assembly mechanisms can vary significantly among different microbial functional groups^24^. However, microbial succession has rarely been studied based on functional trait compositions^14,25^, particularly in response to human-induced environmental changes, such as N addition. Through intensive agriculture, humans have greatly accelerated global N deposition over past centuries^26,27^. Increasing N addition through the application of fertilizers or atmospheric deposition greatly enhances global plant productivity but also dramatically impacts the above- and below-ground biodiversity in terrestrial ecosystems^27–29^. Excessive N input has multiple effects on soil microbial community composition, diversity, activity and function^30–32^. Long-term fertilization is expected to have dramatic and persistent impacts on soil characteristics and microbial community structure^32–35^. For instance, an altered soil C content, C:N ratio, and pH were found to be major factors contributing to variations in microbial communities under fertilization^36–38^, suggesting the importance of deterministic processes (e.g. abiotic filtering effects of environmental selection). In contrast, examining both deterministic and stochastic changes revealed that the effect of N fertilizer was largely exhibited through stochastic processes rather than deterministic ones^39,40^, indicating that complicated assembly mechanisms underlie fertilization effects. A meta-analysis based on 107 datasets from 64 long-term experiments (ranging from 5 to 130 years in length, average 37 years) worldwide revealed that mineral fertilizer application led to a 15% increase in microbial biomass relative to unfertilized soils^30^. However, it remains unknown how long-term anthropogenic fertilization affects the ecological succession of microbial functional guilds and its underlying mechanisms, which is critical for predicting the consequences of increasing anthropogenically derived N addition on terrestrial ecosystems.

The Park Grass Experiment (PGE) on permanent grassland at the Rothamsted Experimental Station, United Kingdom, is the oldest grassland experiment in the world^41^. Since 1856, when the experiment was initiated, plots have received different combinations and amounts of inorganic N, P, and K fertilizers and lime, while the “Nil” control plots have never received fertilizer. Thus, changes in soil geochemical characteristics and soil biota indicate long-term responses to anthropogenic fertilization^41–43^. Archived soil samples from the PGE provide a unique opportunity to address the long-term succession and the underlying assembly mechanisms of soil microbial communities, as influenced by fertilization. In this study, we examined archived soil samples from 1870 to 2008 from contrasting plots with inorganic fertilization and the corresponding “Nil” samples as controls. Soil microbial communities were analyzed using a GeoChip-based metagenomic technology^44,45^ to answer the following questions: (i) What are the succession patterns of soil microbial communities over a century of fertilization, (ii) how does the successional rate of microbial communities change with time, and (iii) how is the century long succession of soil microbial communities controlled by deterministic and stochastic processes? Our results indicate that intensive N addition greatly decreases microbial temporal turnover accompanied by a decrease of stochastic community assembly.

## Results

### Overall pattern of microbial functional communities

An average of 25,842 and 29,541 functional genes were detected in the control and fertilized soils, respectively (Fig. S1). The number of genes overlapping between pairwise samples significantly decreased with time in both control (*p* = 0.048) and fertilized soils (*p* < 0.0001). The proportion of overlapped genes under fertilization was much higher than that in the control. Accordingly, the number of unique genes in the fertilized samples was relatively low. Detrended correspondence analysis (DCA) was performed to visualize the overall patterns of microbial functional communities (Fig. 1a). The control and fertilized samples were clearly separated, indicating dissimilar microbial functional communities between control and long-term inorganic fertilized plots. The microbial communities from fertilized soils were more tightly clustered than those from control soil. Additionally, the dispersion of samples around the group centroid^46^ was quantified. Long-term fertilization led to microbial community convergence (*p* < 0.01) (Fig. 1b). Moreover, three complementary nonparametric multivariate statistical tests (Adonis, ANOSIM and MRPP) further revealed that the overall microbial functional structures were significantly different (*p* < 0.01) between the fertilized and control plots over the century (Table 1). These results indicated that long-term inorganic fertilization significantly altered the functional composition and structure of soil microbial communities.

**Fig. 1.**
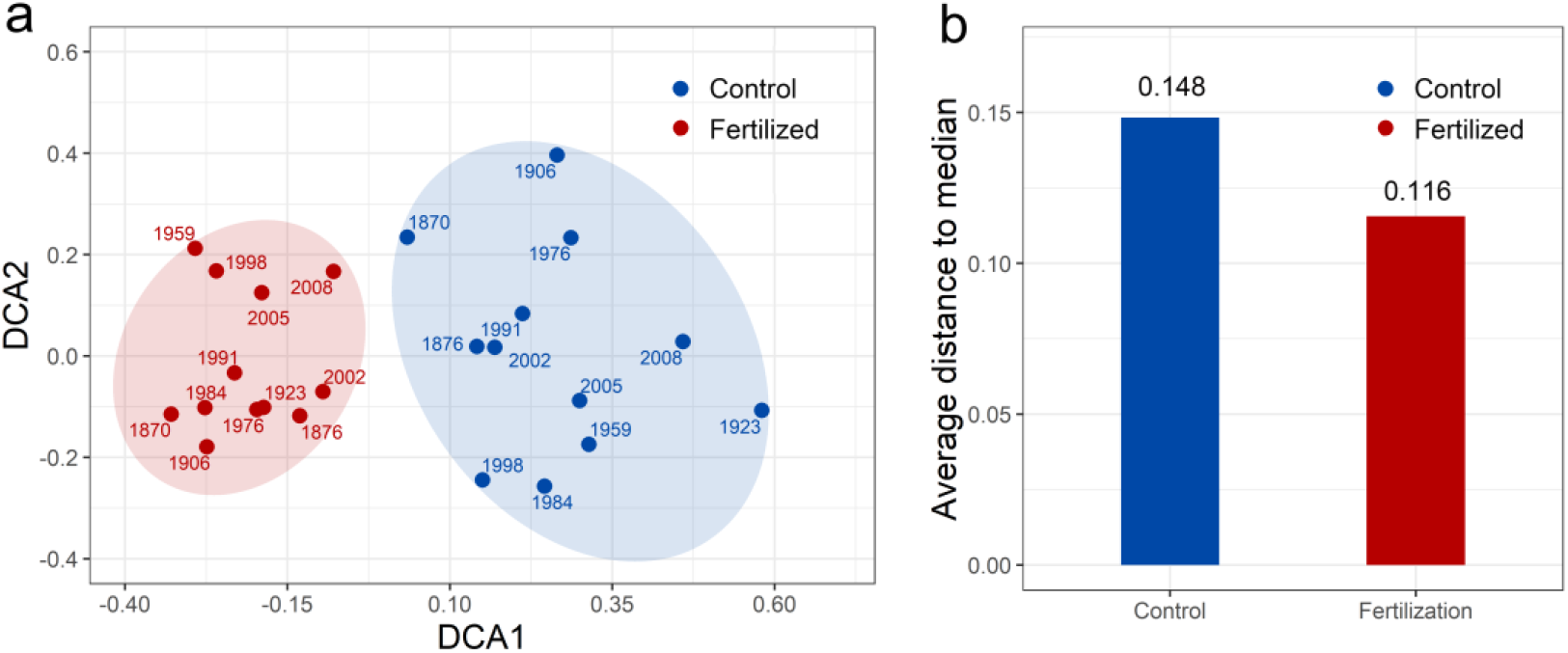
Effects of century-long fertilization on the distributions of soil microbial communities. (a) Overall pattern of microbial functional communities from 1870 to 2008 by Detrended Correspondence Analysis (DCA). Numbers beside the dots indicate sampling years. (b) Dispersion of soil microbial communities in control and fertilized samples.

**Table 1.** Significance tests of the effects of inorganic fertilization on the microbial functional community structure across century long-term with three different statistical approaches.

| Data sets | ANOSIM |  | Adonis |  | MRPP |  |
| --- | --- | --- | --- | --- | --- | --- |
| | R | *p* | F | *p* | $\delta$ | *p* |
| All functional genes | 0.641 | 0.001 | 9.397 | 0.001 | 0.197 | 0.001 |
| C cycling | 0.615 | 0.001 | 8.639 | 0.001 | 0.199 | 0.001 |
| N cycling | 0.658 | 0.001 | 9.687 | 0.001 | 0.195 | 0.001 |
| Phosphorus | 0.703 | 0.001 | 10.582 | 0.001 | 0.212 | 0.001 |
| Sulfur | 0.670 | 0.001 | 9.630 | 0.001 | 0.214 | 0.001 |

All three tests are non-parametric multivariate analyses based on Bray–Curtis dissimilarities among samples, including analysis of similarity (ANOSIM), the permutational multivariate analysis of variance (Adonis), and multiple response permutation procedure (MRPP).

### Microbial temporal turnover under long-term fertilization

Microbial community succession patterns were estimated by linear regression between double log-transformed community similarity (1 – Bray-Curtis distance) and time intervals (Fig. 2a). As expected, significant TDRs were observed for both control (slope = 0.028, *p* = 0.005) and fertilized soils (slope = 0.018, *p* = 0.001). Long-term fertilization significantly decreased microbial temporal turnover rates as determined by a pairwise t-test based on bootstrapping (*p* < 0.0001), which indicated greater community similarity in fertilized soils than in control soil. The effect of DNA degradation with storage time was normalized by applying the same molar quantity of DNA to each GeoChip array^47^, regardless of extraction yield. Whether or not DNA recovery was considered, temporal turnover decreased under long-term fertilization (Fig. S2).

**Fig. 2.**
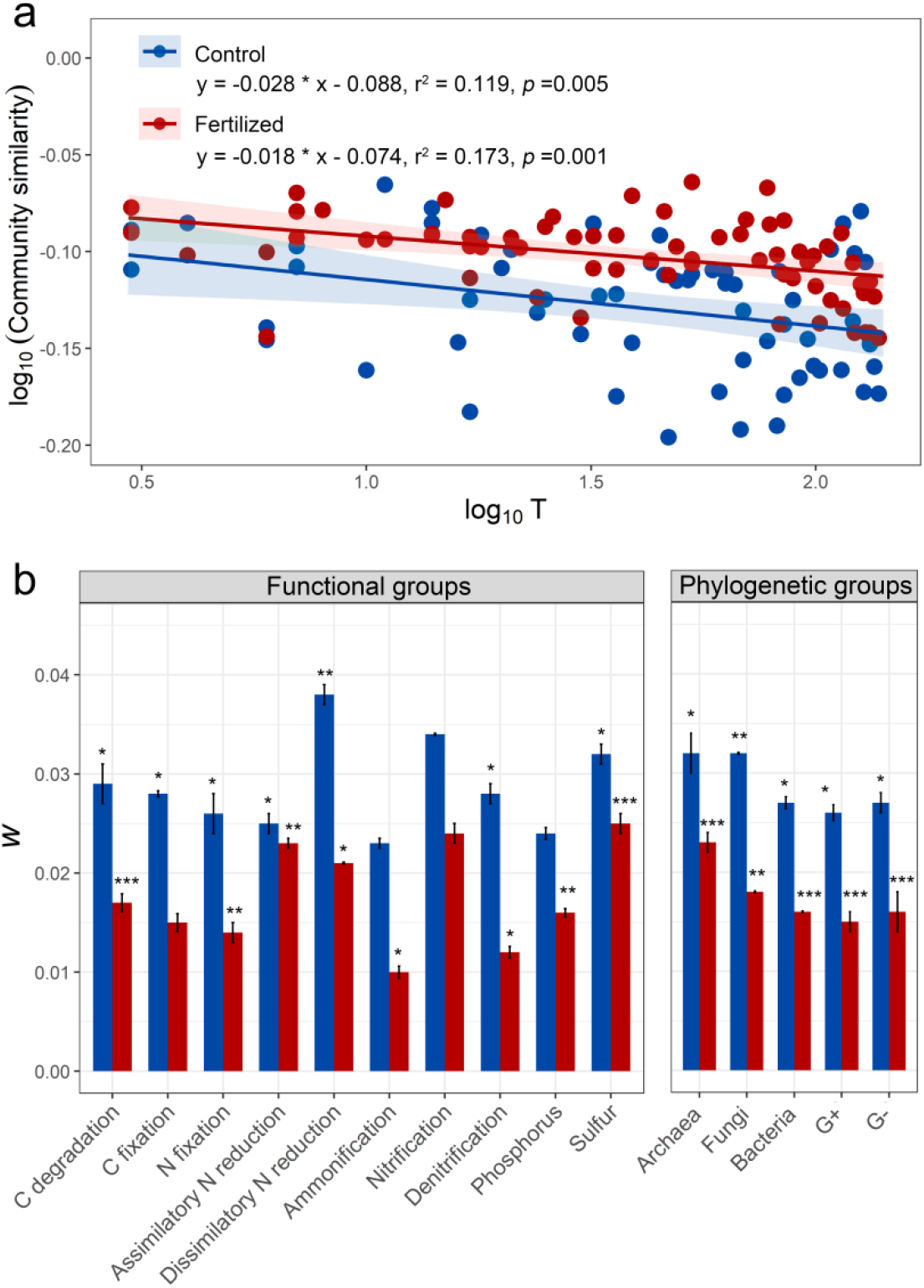
Time-decay relationships (TDR) of soil microbial communities across more than a century. (a) Linear regression between community similarity (*Ss*) and logarithmic time difference (T) of fertilized (red) and control samples (blue), respectively. (b) Temporal turnover (slope of the linear regression) of functional and phylogenetic groups. *<0.05, **<0.01, ***<0.001

The slopes of the TDRs for various functional (C, N, P and S cycling) and phylogenetic (archaea, fungi and bacteria) groups were estimated to obtain additional insights into the temporal changes of different functional and phylogenetic groups in response to fertilization (Fig. 2b). Significant TDRs were found for most functional groups in both control and fertilized soils, with nitrification groups being an exception. Additionally, functional genes involved in ammonification and phosphorus cycling in control soils did not show significant TDRs. Denitrification groups exhibited the greatest difference of turnover rate between fertilized and control samples. Significant TDRs were observed for archaea, fungi and bacteria in both control and fertilized soil samples (*p* < 0.05). In addition, long-term fertilization decreased temporal turnover for all phylogenetic groups (Table S1). The greatest differences were found in fungi (*p* < 0.0001), followed by gram-positive bacteria.

### Time-scale dependence of microbial successional rates

A meta-analysis was conducted to further understand the long-term microbial succession pattern over time periods (Fig. 3, Table S2). Microbial temporal turnovers varied considerably, ranging from 0.001 to 0.31, in different habitats and across different time scales (days to more than a century). A conceptual model with a corresponding mathematical description (rational function) was proposed (Fig. 3a), which is hypothesized to have three phases: In Phase I, *w* increases with time scale (probably from minutes to days or weeks); In Phase II, *w* becomes more or less constant at time scale probably from weeks to years; In Phase III, *w* decreases as the time scale becomes very large, and hence a long tail phase where *w* consistently decreases along time (details in the Methods). All the published TDR slopes and those from our study were tested as a function of their time scales, and the rational function model showed a better fit than other models (Fig. 3b, Table S3). This result supported the above conceptual model of the time-scale dependence of microbial succession rates. Since *w* will decrease and be less and less detectable in the last phase, an interesting question is how long is long enough for time-decay study. This time scale threshold can be defined as the tipping (or threshold) point after which *w* cannot be differentiable from a random pattern. The minimum nonrandom *w* value (*w_t_*) is estimated as 0.0025 and the estimated tipping point is approximately160 years.

**Fig. 3.**
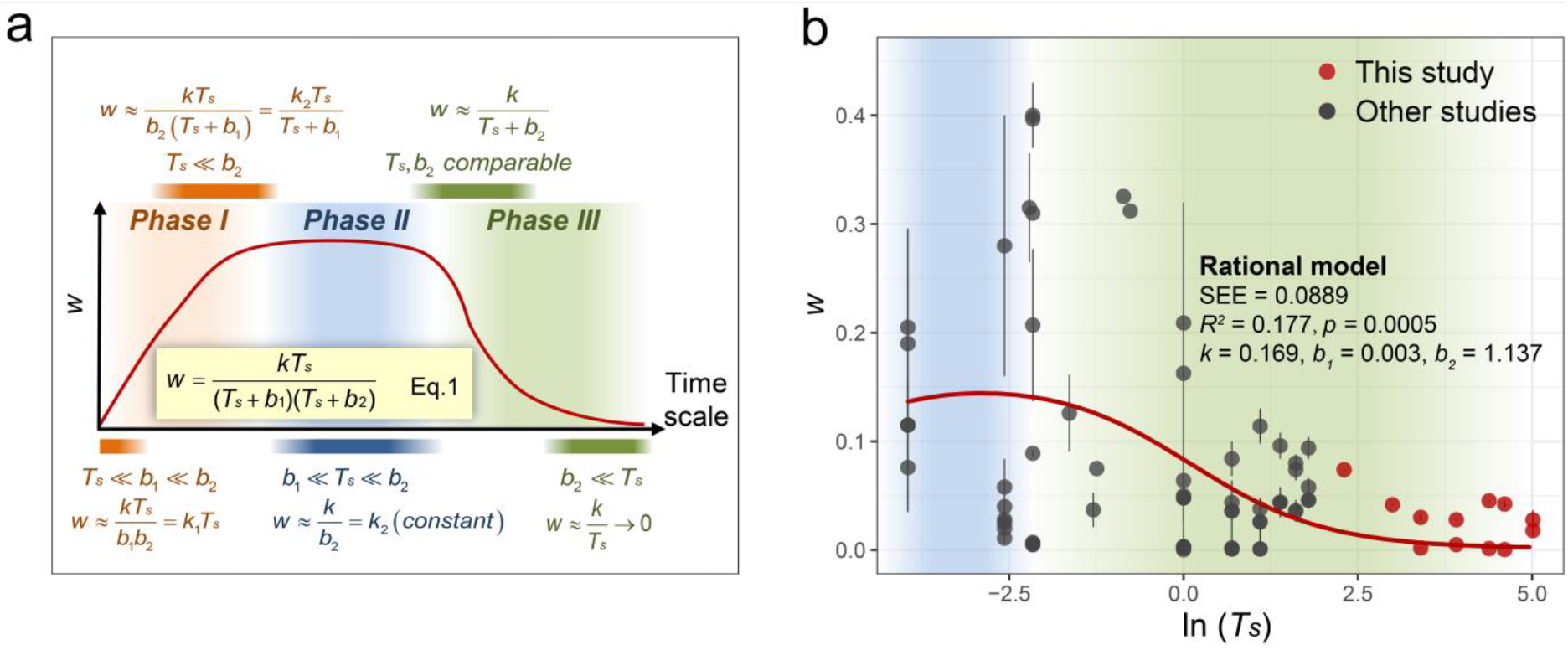
Time-scale dependence of microbial temporal turnover (*w*). (a) Conceptual model with mathematical description for time-scale dependence of microbial temporal turnover. The change in temporal turnover rate (*w*) with time scale (red line) may show different features at three different scales (orange, Phase I; blue, Phase II; green, Phase III). The trend can be described by a rational function model (Eq. 1) that has different approximate formulas at different time scales (phases). *T_s_*, time scale, in year; k, b_1_, b_2_, k_1_, k_2_, constant coefficients. (b) A meta-analysis was conducted to fit the proposed rational function model, as well as other models for comparison (supplementary Table S3). The red line represents the best-fitted rational function model. SEE, standard error of the estimate (details in Table S3). Succession data of microbial communities were obtained from 76 datasets in published studies (supplementary Table S2) and this study. The habitats include air, marine, leaf, surface water, wastewater treatment and soil habitats.

### Ecological stochasticity under long-term fertilization

Null model-based quantification of ecological stochasticity was conducted to test the hypothesis that long-term inorganic fertilization alters the relative importance of stochastic and deterministic processes in driving community succession (Fig. 4). Community assembly was relatively more stochastic in the control plot than in the fertilized plots, with a mean stochasticity ratio (*ST*) of 0.59 (73% *ST*> 0.5) and 0.49 (42% *ST* > 0.5), respectively (paired Wilcoxon test *p* < 0.0001). The temporal succession changed from relatively deterministic (3-year *ST*=0.35~0.45 for control and fertilization) to more stochastic (138-year *ST* = 0.82 for control and 0.68 for fertilization), suggesting that stochastic processes became increasingly important with increasing lengths of community succession. Moreover, fertilization significantly slowed down the increase rate of stochastic processes, resulting in an *ST*-time slope of nearly half that of the control (slope = 0.132 for control and 0.070 for fertilization, bootstrapping test *P* < 0.0001).

**Fig. 4.**
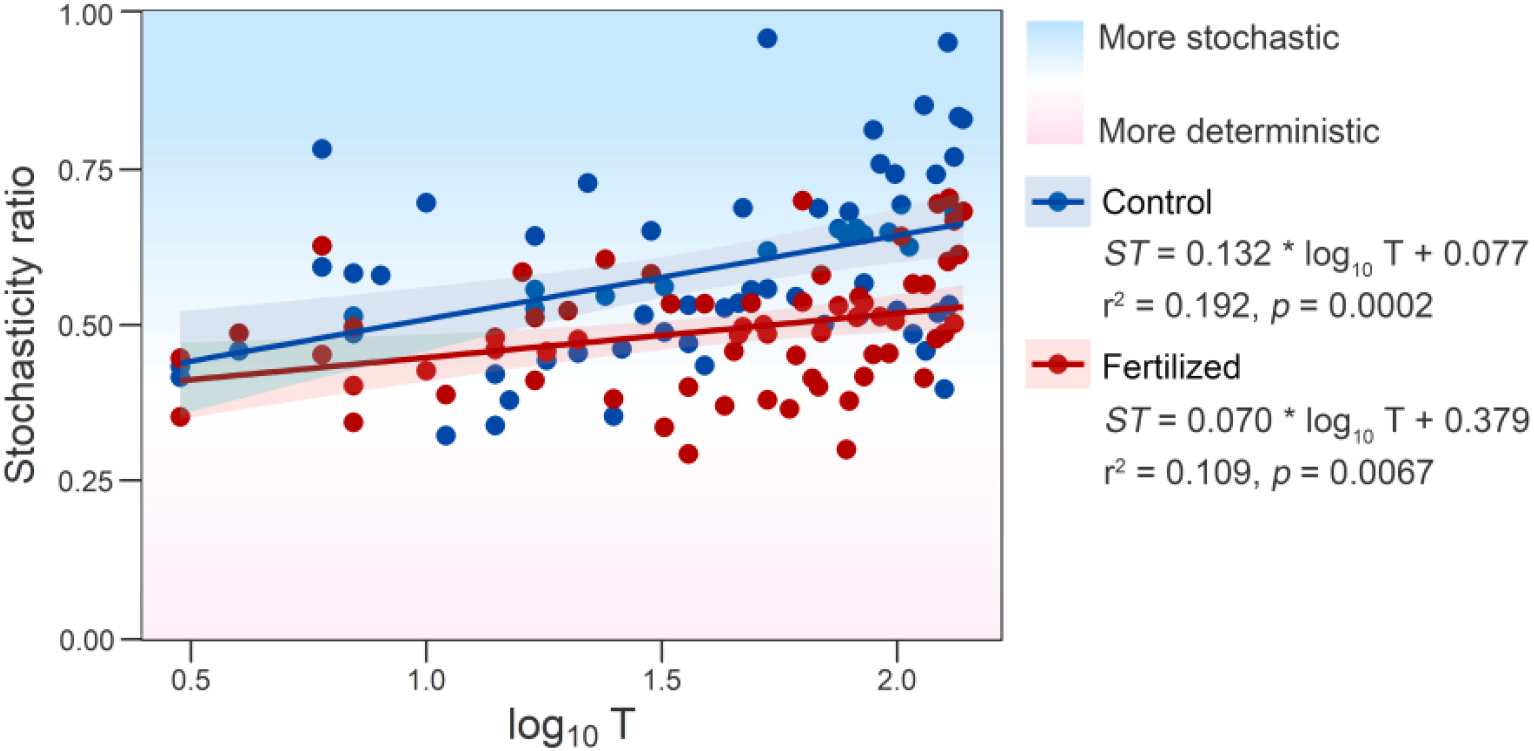
Effect of fertilization on community assembly stochasticity and its temporal change based on null model analysis. The stochasticity ratio was calculated from each pairwise comparison between time points in each plot.

We further compared the effects of fertilization on the ecological stochasticity of different soil functional guilds involved in C and N cycling. As shown in the conceptual diagram (Fig. 5, Table S4), nutrient amendment directly increased soil N availability and indirectly increased soil labile and recalcitrant C pools by enhancing aboveground biomass. Simultaneously, century-long fertilization showed medium-to large-effect-size (Cohen’s *d* = −0.603~ −1.459) decreases in ecological stochasticity in the succession of functional guilds involved in C fixation and degradation; N mineralization, fixation and reduction; and denitrification, coupled with significantly (*p* < 0.0001) decreased TDR slopes. Only *hao* genes were detected from the nitrification group, and the direct addition of sodium nitrate as fertilizer only slightly (Cohen’s *d* = −0.398) decreased the stochasticity of succession in this group, corresponding to a nonsignificant change in the TDR slope.

**Fig. 5.**
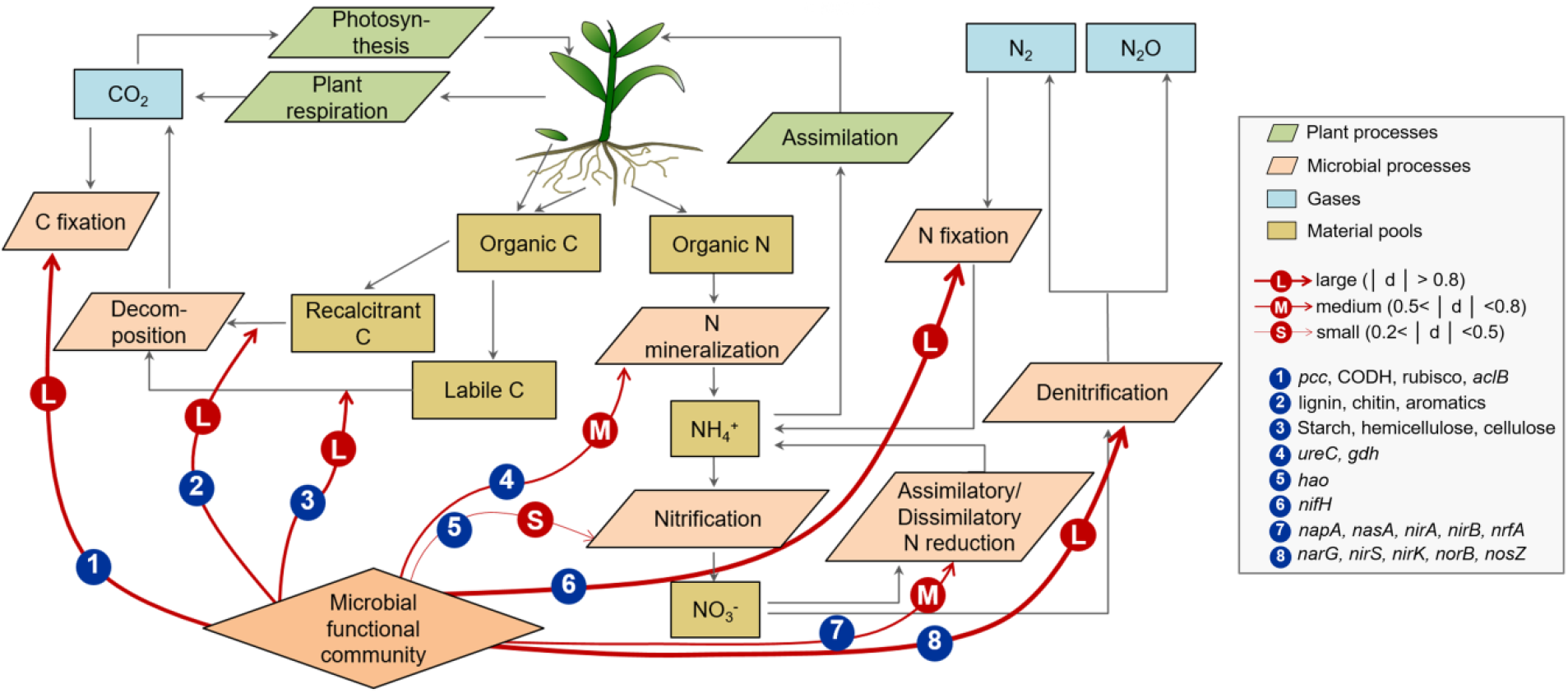
A conceptual diagram of the impact of long-term fertilization on the assembly of microbial functional communities and its potential impacts on microbial functions and ecological variables. Red circles denote impacts on the underlying ecological stochasticity, based on the standardized effect size of fertilization on the stochasticity ratio of each gene group, all of which decreased (negative Cohen’s *d* values) in response to fertilization (supplementary Table S4). Labels of “L”, “M”, and “S” in red circles and the widths of the red lines are proportional to absolute values of Cohen’s *d* and indicate that the decrease in stochasticity was large (|*d*| >0.8), medium (0.5< |*d*| <0.8), or small (0.2< |*d*| <0.5) in effect size. Black arrows indicate material flows. Material pools are represented by yellow rectangles, gases by blue rectangles, microbial processes by orange parallelograms, and plant processes by green parallelograms.

## Discussion

Assembly mechanisms shaping community structure have recently been of a great interest^48–51^, but the drivers controlling ecological succession in response to environmental perturbations, especially on long-term time scales are poorly understood^52,53^. Some stochastic processes, such as dispersal, are likely to be particularly relevant to understanding the temporal variations in β-diversity and succession over a long period of time^54,55^. Here, our null model indicated that the succession of microbial functional structure was increasingly driven by stochastic processes as time proceeded. This result is in accordance with the increasing importance of stochastic succession previously observed in taxonomic/phylogenetic turnovers of microbial communities in resource-rich or low-stress environments^12–14^. This pattern of increasing stochasticity might be due to the cumulative effects of neutral dispersal and random drift over time and/or decrease in cumulative deterministic linkages under unidirectional selection across a long time scale^56^. Furthermore, we found that fertilization reduced the stochasticity of functional structure turnover across the century, corresponding to previous findings that high determinism drove phylogenetic turnovers under selective pressures^13,15,17^. Similarly, we recently reported that a 6-year warming significantly decreased the stochasticity of phylogenetic succession in grassland bacterial communities^8^. These results together suggest that human activities and consequent climate change could impose relatively directional selective pressure(s) and thus change long-term turnover of community composition and functioning towards a more deterministic direction.

Our results further indicated that variations in TDRs of different microbial functional groups were strongly associated with a reduction in stochastic assembly. Inorganic fertilization may exert different selective effects on the regional species pool and specific species^57^ through changes in soil C and N stoichiometric characteristics^58^. Varied niche selection could decrease the importance of stochastic factors in affecting microbial functional community succession to different extents. As in this study, nitrate fertilization increased the concentration of a specific substrate for denitrifiers and stimulated plant growth and thus increased specific C and organic N plant inputs (Fig. S3). Hence, large effects derived from the decrease in stochastic assembly were observed for functional groups involved in C degradation and fixation, N fixation, and denitrification. Correspondingly, of the genes involved in N cycling, nitrification genes were the least impacted by the changing importance of deterministic factors. This behavior may be because the nitrifiers are less abundant and more specialized than other taxa involved in N cycling. There are relatively few ammonia oxidizing bacteria (and archaea) and nitrite oxidizers, and they are probably less resilient, so only those adapted to the PGE plot conditions can survive, resulting in limited potential for stochastic or deterministic divergence. Second, fertilization could also indirectly enhance the selection of host-associated microbial communities through changing plant diversity and biomass^59^. Considerable species loss following N enrichment is observed across terrestrial herbaceous ecosystems^60^. Plant species diversity significantly decreased with long-term fertilization in the PGE^61^, which may result in a more specialized habitat and is another important selection process^62,63^. This trend could explain why fertilization showed large to medium effects on the guilds related to plant C and organic N transformation and root-associated N fixation.

In addition, despite this is the first study to understand the microbial succession and assembly mechanism across more than a century long time, there are some limitations. First, due to historical issues, replicate plots are not available for our study from this unique experiment site. The fertilization is most likely the major reason of the significant difference between the two plots, since the adjacent plots are highly similar in other environmental properties and the significant trends were observed from multiple-time observations across a century. Nevertheless, without replicate plots, it is difficult to parse out contribution of other factors (e.g. soil heterogeneity) to the between-plot difference. Second, since very limited amount of soil samples can be obtained across such long time, the lack of complete edaphic properties constrains us to tease apart how stochasticity may manifest via dispersal or via environmental filtering. Another issue is that the impact of using archived soil on genomic analysis. The impact of archiving on microbial community DNA includes not only degradation in DNA quantity over time, but also rapid and long-term changes in microbial community composition during air-drying and storage. In this case, the DNA that remains in the air-dried soil definitely could not fully represent what are in the original soil. Indeed, we could not assess the magnitude of the storage-induced biases or how these biases may affect the temporal changes of biodiversity. Since archived soil is the only option here, thus we mainly focus on the comparison of the relative changes between treatment and control. We believe that the findings of this study are robust because biases from above reasons can be amended by relative changes comparison^45,48^.

## Conclusions

Our study demonstrates that intensive N applications resulted in decelerating the temporal turnover of soil functional communities, underpinning the differing influences of assembly mechanisms on century long microbial succession. The importance of stochastic assembly differed for soil functional guilds (C, N, P and S) with century long fertilization. These findings have important implications for predicting the ecological consequences of anthropogenically derived N addition at the century scale. The deterministic factors influencing community assembly in fertilized soils may reflect a community with low functional redundancy and hence possibly high susceptibility to disturbances. Terrestrial microbial communities would be much more convergent with increasing N load. Because N addition reduced stochasticity over time, the communities could converge more quickly to a community state with less stochasticity. This effect should be considered when predicting and protecting ecosystem services of soil microbial communities and informing sustainable agricultural practices. In addition, our study demonstrates the value of archived soils for microbial ecology studies employing metagenomic techniques^64^. The availability of archived soil samples makes it possible to track long-term (over a century) impacts of fertilization on grassland ecosystems and to better understand the succession of microbial communities in response to long-term anthropogenic activities across more than a century. It should be noted that our estimation of ecological stochasticity was based on null model analysis and thus cannot avoid interference from randomness due to some deterministic processes (e.g., environmental stochasticity), statistical uncertainties, and inherently high variation in molecular methods^65,66^. Thus, the values should be viewed as statistically approximate and are better used for relative comparison^48^. However, the long-term average trends, particularly the comparison of average trends between the contrasting treatments, are informative and provide novel and important insights.

## Acknowledgements

This study was supported by the National Science Foundation of China (41430856, 41622104), the UK Biotechnology and Biological Sciences Research Council through an Underwood Fellowship to L.W., the Rothamsted Research’s Institute Strategic Programme – Soil to Nutrition (BBS/E/C/000I0310), the Long-Term Experiments National Capability (BBS/E/C/000J0300), the U.S. National Science Foundation MacroSystems Biology program under contract (NSF EF-1065844), Strategic Priority Research Program (B) of the Chinese Academy of Sciences (XDB15010100) and the Youth Innovation Promotion Association of Chinese Academy of Sciences (grant 2016284).

## Author’s contributions

All authors contributed intellectual input and assistance to this study and manuscript preparation. J.Z. developed the original conceptual idea. Y.L., D.N., N.Z., L.W., I.C., D.N., S.M., J.S., P.H., B.S. and Z.L. contributed reagents and data analysis. L.W., and Z.L. did GeoChip analysis. Y.L., D.N. and J. Z. wrote the paper with help from S.M., P.H., I.C. and J.S.

## Data accessibility

GeoChip data are available at Figshare, DOI: 10.6084/m9.figshare.7418504. All other relevant data are available from the corresponding author upon request.

## Supplementary Materials

Supplementary Tables S1-S4

Supplementary Figures S1-S3

## Methods

### Study sites and sample collection

To understand the impacts of long-term inorganic fertilization on soil microbial functional community structure and succession, archived soil samples were taken from Plot 3 and Plot 14/2 of Park Grass, dating back to 1870, 1876, 1906, 1923, 1959, 1976, 1984, 1991, 1998, 2002, 2005 and 2008 for plot 3 and 1870, 1876, 1906, 1923, 1939, 1959, 1984, 1991, 1998, 2002, 2005 and 2008 for plot 14/2. Plot 3 had not received fertilizer since 1856 (control). Plot 14/2 was fertilized with 96 kg N ha^−1^ y^−1^ as sodium nitrate in spring since 1856 with additional minerals (35 kg P ha^−1^ y^−1^ as triple superphosphate, 225 kg K ha^−1^ y^−1^ as potassium sulfate, 15 kg Na ha^−1^ y^−1^ as sodium sulfate, and 10 kg Mg ha^−1^ y^−1^ as magnesium sulfate, and no lime was applied). In total, there are 24 archived soil samples, namely, 12 control samples and 12 fertilized samples. Soils were sampled from 0-10 cm in depth, air-dried and archived in glass bottles. The soil pH was approximately 5 to 6 in both plots, measured in water. Soil C and N were measured by a LECO analyzer, and soils were extracted for Olsen P measurements by colorimetry. The mean annual yields of dry matter were 2.9 t ha^−1^ and 6.7 t ha^−1^ for Plot 3 and Plot 14/2, respectively. Detailed information on the PGE can be found at http://www.rothamsted.ac.uk.

### DNA extraction and GeoChip hybridization

The archived soil samples were preserved by air-drying, and some were stored for more than 150 years. A previous study from the Rothamsted Broadbalk experiment suggested that air-drying of the soil may have preserved DNA in samples stored more than 150 years through desiccation^67^. Genomic DNA was extracted from archived soil samples^68^. Despite air-drying and long-term storage, high-quality DNA was recovered from the soils archived for more than 150 years (260/280 and 260/230 > 1.8). Direct labeling with Cy-3 using random primers and the Klenow fragment of DNA polymerase I was applied to 1 μg of metagenomic DNA^47^. Labeled DNA was then dried in a SpeedVac (45 °C, 45 min; ThermoSavant). GeoChip 4.0 was used to dissect the microbial community functional structure. Hybridization, imaging, and data preprocessing were performed as previously described^44^.

### Time-decay relationships and time-scale dependence of turnover

Time decay relationships (TDRs) were calculated as follows: log_10_(*S_s_*) = constant - *w* log_10_(*T*), where *S_s_* is the pairwise similarity in community composition (1–Bray-Curtis distance), *T* is the time interval, and *w* is a measure of the rate of species turnover with time (temporal turnover). Comparisons of *w* values between control and fertilized samples and *p* values were achieved by bootstrapping (999 times), followed by a pairwise *t*-test.

A conceptual model with a corresponding mathematical description (rational function) was proposed in this study (Fig. 3a). In the early phase, the time difference is too short (e.g., within several seconds) for the community to show obvious change, especially when a diverse microbial community can be resistant to change, i.e., when time interval is close to zero, the TDR slope *w* is close to zero. Then, on a short time scale (probably from minutes to days or weeks), community changes become detectable; thus, the *w* should increase from zero to a certain level. In the rational function model, since *T_s_*<<b_1_<<b_2_ or *T_s_*<<b_2_ in this phase, *w* should increase as a linear function or a Michaelis-Menten-Equation-akin function of time scale (Phase I). In the stable phase, *w* becomes relatively stable probably over periods from weeks to years. In the rational function model, since b_1_<<*T_s_*<<b_2_, *w* becomes a constant and remains stable in this stable phase (Phase II). There is no reason that community similarity will keep decreasing at the same rate forever. The slope *w* should decrease as the time scale becomes very large. Thus, a tail phase where *w* consistently decreases along time and gradually becomes close to zero is proposed. In the rational function model, since *T_s_*>>b_1_ or *T_s_*>>b_2_>>b_1_ in this phase, *w* decreases as a reciprocal function of time scale in this tail phase (Phase III). The early phase (Phase I) is proposed based on reasonable inference, but current studies have not yet covered these effects on such a short time scale. The stable phase (Phase II) has been demonstrated by a previous meta-analysis showing that temporal variability in both within- and between-community diversity was consistent among microbial communities from similar environments^10^. The tail phase (Phase III) can be tested with only long-term datasets, which is now enabled by our century-scale study.

The temporal turnover rates (TDR slope *w* values) were collected from previously published microbial studies (Table S2), where the time scale ranged from days to years. From our century-long study, temporal turnover rates were calculated using sub-dataset across decades in addition to the overall temporal turnover rate across 138 years. All these *w* values and their corresponding time scale (*T_s_*) were used to fit a rational function model (Eq. 1 in Fig. 3a) corresponding to the conceptual model for scale dependence of temporal turnover proposed in this study. The model fitting was compared with fitting using six other models, including logarithmic, power law, reciprocal, quadratic, cubic, and linear models (Table S3). Linear and other nonlinear models were implemented by using the functions “lm” and “nls” in the R package “stats”. Since coefficient of determination (R^2^) can be inadequate for nonlinear models^69^, model fitting was compared by standard error of the estimate (SEE)^70^, accuracy (*χ_a_*) and precision (*ρ*) coefficients derived from concordance correlation coefficient^71^, in addition to R^2^. The significance of each model was calculated using a permutational test in which the time-scale values were permutated 2000 times, and the observed SEE was compared with those of permutated data to calculate the *p* value.

The time-scale threshold, after which temporal turnover cannot be differentiable from random patterns, was calculated as the time scale for a minimum nonrandom *w* value according to the rational function model (Eq.1 in Fig. 3a). The minimum nonrandom *w* value is calculated as the 95% quantile of *w* values from null communities randomized using the null model described below. Considering the uncertainty of coefficient values (*k, b_1_, b_2_*), *w*-vs-*T_s_* data points were randomly drawn (bootstrapping) 1000 times to calculate the coefficients of the rational function model. Then, 1000 estimated time-scale threshold values were calculated to obtain the 95% quantile as a conservative estimation of the threshold, after which the *w* could cease to be meaningful (*p* <0.05).

### Estimating stochasticity of community assembly

Null models are an essential tool for assessing the issue of multiple assembly processes by mimicking the consequences of random processes^72,73^. In a previous study^14^, we assessed the relative importance of stochastic processes with the *ST*, i.e., the ratio of mean expected similarity in the null model 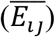 to observed similarity (*C_ij_*), when deterministic factors generally drove community structure to be more similar than the null expectation 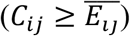. In the present study, *ST* is calculated with some modifications as below to be more general^8^.

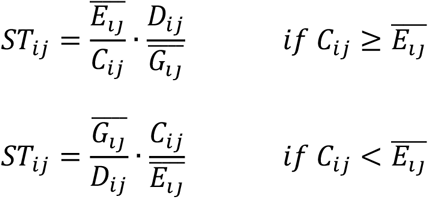

where *C_ij_* is the observed similarity between samples *i* and *j, D_ij_* is the dissimilarity, and *D_ij_*=1-*C_ij_*. *E_ij_* is an expected similarity between null samples *i* and *j*, while *G_ij_* is an expected dissimilarity in the null model. Thus, *ST_ij_* ranges from 0 to 1, where the minimum value is zero when extremely deterministic assembly drives communities completely the same or totally different (*C_ij_* = 0 or *D_ij_*= 0), and the maximum value is one when completely stochastic assembly makes communities the same as null expectation (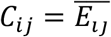 and 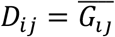). Considering that relative abundances carry information useful for understanding ecological processes^74,75^, the similarity/dissimilarity was measured by the abundance-weighted Bray-Curtis index. The null communities are generated by randomizing the observed community structure 1,000 times based on a null model algorithm described previously^75^. Since we focus on mechanisms underlying temporal succession rather than spatial turnover, the null hypothesis is stochastic succession over time, i.e., the detection or non detection of an individual of a certain species at a time point is due to stochastic processes (e.g., neutral dispersal from a metacommunity, stochastic birth/death, and random diversification) without any determinism (e.g., environmental filtering, competition, mutualism, and evolution under natural selection). Accordingly, all samples from the same plot at different time points are assumed to be from the same species pool across time in our null model algorithm. During randomization, the richness (observed gene numbers) and total gene abundance in each sample are fixed as observed, while the occurrence probability of each gene is proportional to its observed occurrence frequency, and the abundance of each gene is based on random draws of individuals with probability proportional to its relative abundance in the regional species pool. Then, 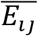 is calculated as the average null expected similarity between samples *i* and *j* and 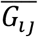 as the average null expected dissimilarity. *ST_ij_* was calculated between every two time points for each plot. Then, the differences in *ST_ij_* values between the control and fertilized plots were tested with a paired *t*-test. The relationship between the pairwise *ST_ij_* and the time interval in each plot was fit to a linear model, and the significance of each model and the slope difference between control and fertilized plots were tested in the same way as described for the time-decay model.

Furthermore, *ST_ij_* was also separately calculated for each of the major C and N cycling functional gene groups, considering that ecological stochasticity and its response to fertilization could be different in different guilds. Subcommunities were extracted for each gene group, including the nitrification group, the denitrification group, the labile C degradation group, etc. Then, the *ST_ij_* of each gene group was calculated using the same method described above. To evaluate the effect of fertilization on ecological stochasticity, Cohen’s *d* (paired per time point)^76^ was calculated between the *ST_ij_* values in the control and fertilized plots for each gene group. The standardized effects are large, medium, small or negligible when |*d*| > 0.8, 0.5 < |*d*| < 0.8, 0.2 < |*d*| < 0.5, or |*d*| < 0.2,, respectively, and *d* values can be positive or negative to reflect increases or decreases in the standardized effect size of fertilization on the stochasticity ratio.

### Other statistical analyses

Detrended correspondence analysis (DCA) were used to determine the overall pattern of microbial functional communities. The microbial temporal patterns under fertilized and control plots were also determined by non-metric multidimensional scaling ordination based on the Bray–Curtis dissimilarity. Three different non-parametric multivariate statistical tests (analysis of similarity (ANOSIM), nonparametric multivariate analysis of variance (Adonis), and multi-response permutation procedure (MRPP)) were used to test the differences in soil microbial communities under fertilization and control treatments. The ANOSIM statistic R is based on the difference of mean ranks between groups and within groups and is calculated as follows:

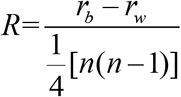

where *r_b_* is the mean rank of between-group dissimilarities, *r_w_* is the mean rank of within-group dissimilarities, and n is the total number. According to the ranks of the dissimilarities, statistic *R* ranges from −1 to +1. Ecological communities rarely achieve an R value <0. Thus, R ≈ 0 indicates no difference among groups, and R > 0 indicates that groups differ in community composition.

## References

1 Michaud, J., Schoenly, K. G. & Moreau, G., Rewriting ecological succession history: Did carrion ecologists get there first? Q REV BIOL 90 45 (2015).

2 Falk, M. W., Seshan, H., Dosoretz, C. & Wuertz, S., Partial bioaugmentation to remove 3-chloroaniline slows bacterial species turnover rate in bioreactors. WATER RES 47 7109 (2013).

3 Jiao, S. et al., Distinct succession patterns of abundant and rare bacteria in temporal microcosms with pollutants. ENVIRON POLLUT 225 497 (2017).

4 Kim, T., Jeong, J., Wells, G. F. & Park, H., General and rare bacterial taxa demonstrating different temporal dynamic patterns in an activated sludge bioreactor. APPL MICROBIOL BIOT 97 1755 (2013).

5 Pla-Rabes, S., Flower, R. J., Shilland, E. M. & Kreiser, A. M., Assessing microbial diversity using recent lake sediments and estimations of spatio-temporal diversity. J BIOGEOGR 38 2033 (2011).

6 Redford, A. J. & Fierer, N., Bacterial succession on the leaf surface: A novel system for studying successional dynamics. MICROB ECOL 58 189 (2009).

7 van der Gast, C. J., Ager, D. & Lilley, A. K., Temporal scaling of bacterial taxa is influenced by both stochastic and deterministic ecological factors. ENVIRON MICROBIOL 10 1411 (2008).

8 Guo, X. et al., Climate warming leads to divergent succession of grassland microbial communities. NAT CLIM CHANGE 8 813 (2018).

9 Liang, Y. et al., Long-term soil transplant simulating climate change with latitude significantly alters microbial temporal turnover. The ISME Journal 9 2561 (2015).

10 Shade, A., Caporaso, J. G., Handelsman, J., Knight, R. & Fierer, N., A meta-analysis of changes in bacterial and archaeal communities with time. The ISME Journal 7 1493 (2013).

11 Xiong, J. et al., Evidence of bacterioplankton community adaptation in response to long-term mariculture disturbance. SCI REP-UK 5 15274 (2015).

12 Ofiteru, I. D. et al., Combined niche and neutral effects in a microbial wastewater treatment community. P NATL ACAD SCI USA 107 15345 (2010).

13 Tripathi, B. M. et al., Soil pH mediates the balance between stochastic and deterministic assembly of bacteria. The ISME Journal 12 1072 (2018).

14 Zhou, J. et al., Stochasticity, succession, and environmental perturbations in a fluidic ecosystem. P NATL ACAD SCI USA 111 E836 (2014).

15 Campisano, A. et al., Temperature drives the assembly of endophytic communities’ seasonal succession. ENVIRON MICROBIOL 19 3353 (2017).

16 Lin, Q. et al., Temperature regulates deterministic processes and the succession of microbial interactions in anaerobic digestion process. WATER RES 123 134 (2017).

17 Zhalnina, K. et al., Dynamic root exudate chemistry and microbial substrate preferences drive patterns in rhizosphere microbial community assembly. NAT MICROBIOL 3 470 (2018).

18 Darcy, J. L. et al., Phosphorus, not nitrogen, limits plants and microbial primary producers following glacial retreat. SCI ADV 4 q942 (2018).

19 Knelman, J. E. et al., Nutrient addition dramatically accelerates microbial community succession. PLOS ONE 9 (2014).

20 Rivett, D. W. et al., Elevated success of multispecies bacterial invasions impacts community composition during ecological succession. ECOL LETT 21 516 (2018).

21 Bardgett, R. D., Bowman, W. D., Kaufmann, R. & Schmidt, S. K., A temporal approach to linking aboveground and belowground ecology. Trends in Ecology and Evolution 20 634 (2005).

22 Dini-Andreote, F., Pylro, V. S., Baldrian, P., van Elsas, J. D. & Salles, J. F., Ecological succession reveals potential signatures of marine-terrestrial transition in salt marsh fungal communities. The ISME Journal 10 1984 (2016).

23 Fukami, T., in Annual Review of Ecology Evolution and Systematics, edited by D. J. Futuyma (Annual Reviews, Palo Alto, 2015), Vol. 46, pp. 1.

24 Caruso, T. et al., Stochastic and deterministic processes interact in the assembly of desert microbial communities on a global scale. The ISME Journal 5 1406 (2011).

25 Gutknecht, J. L. M., Field, C. B. & Balser, T. C., Microbial communities and their responses to simulated global change fluctuate greatly over multiple years. GLOBAL CHANGE BIOL 18 2256 (2012).

26 Reay, D. S., Dentener, F., Smith, P., Grace, J. & Feely, R. A., Global nitrogen deposition and carbon sinks. NAT GEOSCI 1 430 (2008).

27 Stevens, C. J., Dise, N. B., Mountford, J. O. & Gowing, D. J., Impact of nitrogen deposition on the species richness of grasslands. SCIENCE 303 1876 (2004).

28 Stevens, C. J., David, T. I. & Storkey, J., Atmospheric nitrogen deposition in terrestrial ecosystems: Its impact on plant communities and consequences across trophic levels. FUNCT ECOL 32 1757 (2018).

29 Zhang, T., Chen, H. Y. H. & Ruan, H., Global negative effects of nitrogen deposition on soil microbes. The ISME Journal 12 1817 (2018).

30 Geisseler, D. & Scow, K. M., Long-term effects of mineral fertilizers on soil microorganisms - A review. Soil Biology and Biochemistry 75 54 (2014).

31 Ramirez, K. S., Lauber, C. L., Knight, R., Bradford, M. A. & Fierer, N., Consistent effects of nitrogen fertilization on soil bacterial communities in contrasting systems. ECOLOGY 91 3463 (2010).

32 Rousk, J., Brookes, P. C. & Baath, E., Fungal and bacterial growth responses to N fertilization and pH in the 150-year ‘Park Grass’ UK grassland experiment. FEMS MICROBIOL ECOL 76 89 (2011).

33 Hallin, S., Jones, C. M., Schloter, M. & Philippot, L., Relationship between N-cycling communities and ecosystem functioning in a 50-year-old fertilization experiment. The ISME Journal 3 597 (2009).

34 Liu, W. et al., Changes in the abundance and structure of bacterial communities under long-term fertilization treatments in a peanut monocropping system. PLANT SOIL 395 415 (2015).

35 Zhou, J. et al., Influence of 34-years of fertilization on bacterial communities in an intensively cultivated black soil in northeast China. Soil Biology and Biochemistry 90 42 (2015).

36 Zhong, W. et al., The effects of mineral fertilizer and organic manure on soil microbial community and diversity. PLANT SOIL 326 511 (2010).

37 Xun, W. et al., Significant alteration of soil bacterial communities and organic carbon decomposition by different long-term fertilization management conditions of extremely low-productivity arable soil in South China. ENVIRON MICROBIOL 18 1907 (2016).

38 Zhou, Z., Wang, C., Zheng, M., Jiang, L. & Luo, Y., Patterns and mechanisms of responses by soil microbial communities to nitrogen addition. Soil Biology and Biochemistry 115 433 (2017).

39 Zhang, X., Johnston, E. R., Liu, W., Li, L. & Han, X., Environmental changes affect the assembly of soil bacterial community primarily by mediating stochastic processes. GLOBAL CHANGE BIOL 22 198 (2016).

40 Zhang, X., Liu, W., Bai, Y., Zhang, G. & Han, X., Nitrogen deposition mediates the effects and importance of chance in changing biodiversity. MOL ECOL 20 429 (2011).

41 Silvertown, J. et al., The Park Grass Experiment 1856-2006: Its contribution to ecology. J ECOL 94 801 (2006).

42 Storkey, J. et al., in Advances in Ecological Research, edited by A. J. Dumbrell, R. L. Kordas and G. Woodward (2016), Vol. 55, pp. 3.

43 Crawley, M. J. et al., Determinants of species richness in the park grass experiment. AM NAT 165 179 (2005).

44 Tu, Q. et al., GeoChip 4: a functional gene-array-based high-throughput environmental technology for microbial community analysis. MOL ECOL RESOUR n/a-n/a (2014).

45 Zhou, J. et al., High-throughput metagenomic technologies for complex microbial community analysis: Open and closed formats. MBIO 6 e2214 (2015).

46 Houseman, G. R., Mittelbach, G. G., Reynolds, H. L. & Gross, K. L., Perturbations alter community convergence, divergence, and formation of multiple community states. ECOLOGY 89 2172 (2008).

47 Wu, L., Liu, X., Schadt, C. W. & Zhou, J., Microarray-based analysis of subnanogram quantities of microbial community DNAs by using whole-community genome amplification. APPL ENVIRON MICROB 72 4931 (2006).

48 Zhou, J. & Ning, D., Stochastic community assembly-Does it matter in microbial ecology? MICROBIOL MOL BIOL R 81 e2 (2017).

49 Chase, J. M., Stochastic community assembly causes higher biodiversity in more productive environments. SCIENCE 328 1388 (2010).

50 Ellwood, M. D. F., Manica, A. & Foster, W. A., Stochastic and deterministic processes jointly structure tropical arthropod communities. ECOL LETT 12 277 (2009).

51 Kraft, N. J. B. et al., Disentangling the drivers of beta diversity along latitudinal and elevational gradients. SCIENCE 333 1755 (2011).

52 Ferrenberg, S. et al., Changes in assembly processes in soil bacterial communities following a wildfire disturbance. The ISME Journal 7 1102 (2013).

53 Chang, C. & HilleRisLambers, J., Integrating succession and community assembly perspectives. F1000 Research 5 (2016).

54 Fierer, N., Nemergut, D., Knight, R. & Craine, J. M., Changes through time: Integrating microorganisms into the study of succession. RES MICROBIOL 161 635 (2010).

55 Langenheder, S., Berga, M., Ostman, O. & Szekely, A. J., Temporal variation of beta-diversity and assembly mechanisms in a bacterial metacommunity. The ISME Journal 6 1107 (2012).

56 Nemergut, D. R. et al., Patterns and processes of microbial community assembly. MICROBIOL MOL BIOL R 77 342 (2013).

57 Francioli, D. et al., Mineral vs. organic amendments: microbial community structure, activity and abundance of agriculturally relevant microbes are driven by long-term fertilization strategies. FRONT MICROBIOL 7 (2016).

58 Jian, S. et al., Soil extracellular enzyme activities, soil carbon and nitrogen storage under nitrogen fertilization: A meta-analysis. Soil Biology and Biochemistry 101 32 (2016).

59 Yuan, X. et al., Plant community and soil chemistry responses to long-term nitrogen inputs drive changes in alpine bacterial communities. ECOLOGY 97 1543 (2016).

60 Clark, C. M., Cleland, E. E., Collins, S. L., Fargione, J. E. & Gough, L., Environmental and plant community determinants of species loss following nitrogen enrichment. ECOL LETT 10 596 (2007).

61 Liang, Y. et al., Over 150 years long-term fertilization alters spatial scaling of microbial biodiversity. MBIO 6 e215 (2015).

62 Robinson, C. J., Bohannan, B. J. M. & Young, V. B., From structure to function: the ecology of host-associated microbial communities. MICROBIOL MOL BIOL R 74 453 (2010).

63 Valyi, K., Mardhiah, U., Rillig, M. C. & Hempel, S., Community assembly and coexistence in communities of arbuscular mycorrhizal fungi. The ISME Journal 10 2341 (2016).

64 Cary, S. C. & Fierer, N., The importance of sample archiving in microbial ecology. NAT REV MICROBIOL 12 789 (2014).

65 Zhou, J. et al., Reproducibility and quantitation of amplicon sequencing-based detection. The ISME Journal 5 1303 (2011).

66 Zhou, J. et al., Random sampling process leads to overestimation of beta-diversity of microbial communities. MBIO 4 (2013).

## References

67 Clark, I. M. & Hirsch, P. R., Survival of bacterial DNA and culturable bacteria in archived soils from the Rothamsted Broadbalk experiment. SOIL BIOL BIOCHEM 40 1090 (2008).

68 Zhou, J., Bruns, M. A. & Tiedje, J. M., DNA recovery from soils of diverse composition. APPL ENVIRON MICROB 62 316 (1996).

69 Spiess, A. N. & Neumeyer, N., An evaluation of R^2^ as an inadequate measure for nonlinear models in pharmacological and biochemical research: a Monte Carlo approach. BMC PHARMACOL 10 6 (2010).

70 Weisberg, S., Applied linear regression, 3rd ed. (John Wiley & Sons, NJ 2005).

71 Lin, L., Hedayat, A.S., Sinha, B., & Yang, M., Statistical methods in assessing agreement: models, issues, and tools. J AM STAT ASSOC 97 257 (2002).

72 Chase, J. M. & Myers, J. A., Disentangling the importance of ecological niches from stochastic processes across scales. PHILOS T R SOC B 366 2351 (2011).

73 Myers, J. A. et al., Beta-diversity in temperate and tropical forests reflects dissimilar mechanisms of community assembly. Ecoloty Letters 16 151 (2013).

74 Anderson, M. J. et al., Navigating the multiple meanings of beta diversity: a roadmap for the practicing ecologist. ECOL LETT 14 19 (2011).

75 Stegen, J. C. et al., Quantifying community assembly processes and identifying features that impose them. ISME J 7 2069 (2013).

76 Cohen, J., Statistical power analysis for the behavioral sciences, 2nd ed. (Lawrence Erlbaum Associates, Hillsdale, NJ, 1988).

