## Supplementary tables and figures for "Century long fertilization reduces stochasticity controlling grassland microbial community succession"

**Dr. Yuting Liang**

[#] These authors contributed equally to this work.

### Supplementary Tables

**Table S1. Slopes of time-decay relationships for various functional and phylogenetic groups under control or long-term fertilization in archived soils.**

| | Control | | | Fertilization | | | T test* | |
|---|---|---|---|---|---|---|---|---|
| | slope | $r^2$ | *p* | slope | $r^2$ | *p* | t | *P* |
| ***Functional groups*** | | | | | | | | |
| C degradation | **0.029** | 0.294 | 0.016 | **0.017** | 0.405 | <0.001 | 29.3 | <0.0001 |
| C fixation | **0.028** | 0.287 | 0.019 | **0.015** | 0.397 | <0.001 | 31.67 | <0.0001 |
| N fixation | **0.026** | 0.287 | 0.019 | **0.014** | 0.341 | 0.005 | 30.22 | <0.0001 |
| Assimilatory N reduction | **0.025** | 0.277 | 0.024 | **0.023** | 0.368 | 0.002 | 4.95 | <0.0001 |
| Dissimilatory N reduction | **0.038** | 0.333 | 0.006 | **0.021** | 0.3 | 0.014 | 32.1 | <0.0001 |
| Ammonification | 0.023 | 0.231 | 0.061 | **0.01** | 0.308 | 0.012 | | |
| Nitrification | 0.034 | 0.162 | 0.192 | 0.024 | 0.146 | 0.241 | | |
| Denitrification | **0.028** | 0.271 | 0.028 | **0.012** | 0.316 | 0.01 | 73.1 | <0.0001 |
| Phosphorus | 0.024 | 0.0216 | 0.082 | **0.016** | 0.351 | 0.004 | | |
| Sulfur | **0.032** | 0.264 | 0.032 | **0.025** | 0.523 | <0.001 | 13.94 | <0.0001 |
| ***Phylogenetic groups*** | | | | | | | | |
| Archaea | **0.032** | 0.304 | 0.013 | **0.023** | 0.438 | <0.001 | 19.57 | <0.0001 |
| Fungi | **0.032** | 0.342 | 0.005 | **0.018** | 0.391 | 0.001 | 33.24 | <0.0001 |
| Bacteria | **0.027** | 0.261 | 0.034 | **0.016** | 0.431 | <0.001 | 25.30 | <0.0001 |
| G+ | **0.026** | 0.265 | 0.031 | **0.015** | 0.434 | <0.001 | 25.70 | <0.0001 |
| G- | **0.027** | 0.258 | 0.036 | **0.016** | 0.428 | <0.001 | 25.75 | <0.0001 |

Significant slopes of the linear regression between microbial dissimilarity (Bray-Curtis distance) and log transformed time were bolded. * The right-most columns test whether the slopes between control and fertilization were significantly different.

**Table S2. List of taxonomic groups and references that reported TDR in microbial communities.**

| Taxonomic group | Ecosystem | Time periods | Approaches | References |
|---|---|---|---|---|
| Bacteria | Soil | 100 days | 16S | Jiao, S. *et al.* 2017[1] |
| Bacteria | Marine | 10 years | ARISA* | Cram, A.J. *et al.* 2014[2] |
| Bacteria | Marine | 80 days, 10 years | 16S | Fuhrman, A.J. *et al.* 2015[3] |
| Bacteria, eukaryote | Sediment | 1 year | 16S, 18S | Zhang, N. *et al*. 2017[4] |
| Bacteria | Shrimp ponds | 42 days | 16S | Xiong, J. *et al.* 2014[5] |
| Bacteria | Soil | 100 days | 16S | Jiao, S. *et al.* 2017[6] |
| Bacteria | Desert | 1 year | 16S | Armstrong, A. *et al.* 2016[7] |
| Bacteria | Human | 1 year | 16S | Stressmann, F.A. *et al.* 2012[8] |
| Bacteria and Archaea | Air, plant, lake, stream, marine, human, soil, wastewater | 1 week to 6 years | 16S | Shade, A. *et al.* 2013[9] |
| Bacteria | Marine | Seasonal to 8 years | 16S | Hatosy, S.M. *et al.* 2013[10] |
| Bacteria | Soil | 6 years | 16S | Liang, Y. *et al.* 2015[11] |
| Bacteria and fungi | Tall-grass prairie | 6 years | 16S, ITS | Guo, X. *et al.* 2018[12] |

*ARISA, Automated Ribosomal Intergenic Spacer Analysis

**Table S3 Multiple models to fit the relationship between temporal turnovers (*w*) and time scale (*T*<sub>s</sub>).**

| Model | Equation | SEE | $\chi_a$ | $\rho$ | $R^2$ | *p* |
|---|---|---|---|---|---|---|
| Rational | $w = kT/[(T_s + b_1)(T_s + b_2)]$ | **0.0889** | **0.798** | **0.429** | **0.177** | 0.0005 |
| Logarithmic | $w = k\ \ln(T_s) + b$ | 0.0897 | 0.695 | 0.405 | 0.164 | 0.0005 |
| Pow law | $w = b{T_s}^a$ | 0.0898 | 0.665 | 0.402 | 0.161 | 0.001 |
| Reciprocal | $w = b + (a/T_s)$ | 0.0946 | 0.491 | 0.263 | 0.069 | 0.022 |
| Quadratic | $w = a{T_s}^2 + bT_s + c$ | 0.0947 | 0.486 | 0.259 | 0.067 | 0.0735 |
| Cubic | $w = a{T_s}^3 + b{T_s}^2 + cT_s + d$ | 0.0935 | 0.551 | 0.300 | 0.090 | 0.089 |
| Linear | $w = aT_s + b$ | 0.0958 | 0.551 | 0.300 | 0.045 | 0.058 |

Bold numbers indicate the best fit in the compared models. SEE, standard error of the estimate[16]; $\chi_a$ and $\rho$ are proposed as components in concordance correlation coefficient ($\rho_c$) to evaluate accuracy and precision, respectively[17]; $R^2$, coefficient of determination; *p* values were calculated by permutational test. Equations are as below.

$$SEE = \sqrt{\frac{\sum_i (w_i - \hat{w}_i)^2}{n-2}} \quad \textbf{Eq. S1}$$

$$\chi_a = \frac{2\sigma_w \sigma_{\hat{w}}}{\sigma_w^2 + \sigma_{\hat{w}}^2 + (\mu_w - \mu_{\hat{w}})^2} \quad \textbf{Eq. S2}$$

$$\rho = \frac{\sigma_{w\hat{w}}}{\sigma_w \sigma_{\hat{w}}} \quad \textbf{Eq. S3}$$

$$\rho_c = \chi_a \rho = \frac{2\sigma_{w\hat{w}}}{\sigma_w^2 + \sigma_{\hat{w}}^2 + (\mu_w - \mu_{\hat{w}})^2} \quad \textbf{Eq. S4}$$

$$R^2 = 1 - \frac{\sum_i (w_i - \hat{w}_i)^2}{\sum_i (w_i - \mu_{\hat{w}})^2} \quad \textbf{Eq. S5}$$

| | |
|---|---|
| $w_i$ | Observed *w* value. |
| $\hat{w}_i$ | Estimated (model predicted) *w* value. |
| $n$ | Number of data points. |
| $\mu_w$ | Mean of observed *w* values |
| $\mu_{\hat{w}}$ | Mean of estimated *w* values |
| $\sigma_w$ | Variance of observed *w* values |
| $\sigma_{\hat{w}}$ | Variance of estimated *w* values |
| $\sigma_{w\hat{w}}$ | Covariance of observed and estimated *w*. |

**Table S4 Cohen's *d* values to evaluate the effect of long-term fertilization on stochastic assembly of microbial functional groups.**

| Functional processes | Functional genes | *d* | Effect size |
|---|---|---|---|
| C fixation | *pcc*, CODH, rubisco, *aclB* | -1.23 | Large |
| Recalcitrant C degradation | lignin, chitin, aromatics | -0.90 | Large |
| Labile C degradation | starch, hemicellulose, cellulose | -1.00 | Large |
| N mineralization | *ureC*, *gdh* | -0.60 | Medium |
| Nitrification | *hao* | -0.40 | Small |
| N fixation | *nifH* | -1.46 | Large |
| Assimilatory/dissimilatory N reduction | *napA*, *nasA*, *nirA*, *nirB*, *nrfA* | -0.62 | Medium |
| Denitrification | *narG*, *nirS*, *nirK*, *norB*, *nosZ* | -1.34 | Large |

References:


1. Jiao, S., Zhang, Z., Yang, F., Lin, Y., Chen, W., & Wei, G. Temporal dynamics of microbial communities in microcosms in response to pollutants. *MOL ECOL* **26** 923 (2017).
2. Cram, J.A., Chow, T.C.E., Sachdeva, R., Needham, D.M., Parada, A.E., Steele, J.A., & Fuhrman, J.A. Seasonal and interannual variability of the marine bacterioplankton community throughout the water column over ten years. *ISME J* **9** 563 (2015).
3. Fuhrman, J.A., Cram, J.A., & Needham, D.M. Marine microbial community dynamics and their ecological interpretation. *NAT REV MICROBIOL* **13** 133 (2015).
4. Zhang, N., Xiao, X., Pei, M., Liu, X., & Liang, Y. Discordant temporal turnovers of sediment bacterial and eukaryotic communities in response to dredging: nonresilience and functional changes. *APPL ENVIRON MICROBIOL* 83 e02526-16 (2017).
5. Xiong, J., Zhu, J., Wang, K. *et al*. The temporal scaling of bacterioplankton composition: high turnover and predictability during shrimp cultivation. *MICROB ECOL* **67** 256 (2014).
6. Jiao, S., Luo, Y., Lu, M. *et al*. Distinct succession patterns of abundant and rare bacteria in temporal microcosms with pollutants. *ENVIRON POLLUT* **225** 497 (2017).
7. Armstrong, A., Valverde, A., Ramond, J.B. *et al*. Temporal dynamics of hot desert microbial communities reveal structural and functional responses to water input. *SCI REP-UK* **6** 34434 (2016).
8. Stressmann, F.A., Rogers, G.B., van der Gast, C.J. *et al*. Long-term cultivation-independent microbial diversity analysis demonstrates that bacterial communities infecting the adult cystic fibrosis lung show stability and resilience. *THORAX* **67** 867 (2012).
9. Shade, A., Caporaso, J.G., Handelsman, J. *et al*. A meta-analysis of changes in bacterial and archaeal communities with time. *ISME J* **7** 1493 (2013).
10. Hatosy, S.M., Martiny, J.B.H., Sachdeva, R. *et al*. Beta diversity of marine bacteria depends on temporal scale. *ECOLOGY* **94** 1898 (2013).
11. Liang, Y., Jiang, Y., Wang, F. *et al*. Long-term soil transplant simulating climate change with latitude significantly alters microbial temporal turnover. *ISME J* **9** 2561 (2015).
12. Guo, X., Feng, J., Shi, Z. *et al*. Climate warming leads to divergent succession of grassland microbial communities. *NAT CLIM CHANGE* **8** 813 (2018).
13. Weisberg, S., *Applied linear regression*, 3rd ed. (John Wiley & Sons, NJ 2005).
14. Lin, L., Hedayat, A.S., Sinha, B., & Yang, M., Statistical methods in assessing agreement: models, issues, and tools. *J AM STAT ASSOC* **97** 257 (2002).

### Supplementary Figures

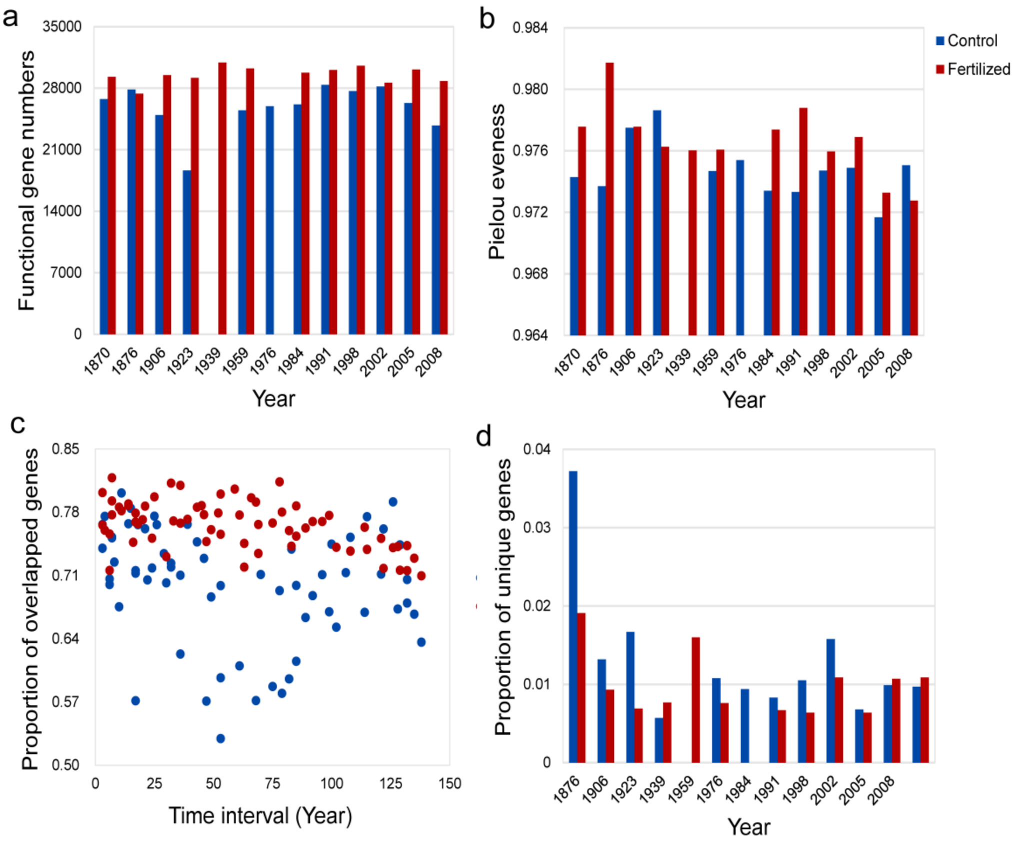


**Fig. S1**. **Effects of long-term fertilization on microbial functional genes**. (a) Functional gene numbers, (b) Pielou evenness, (c) percentage of overlapped genes between pairwise samples, and (d) percentage of unique genes.

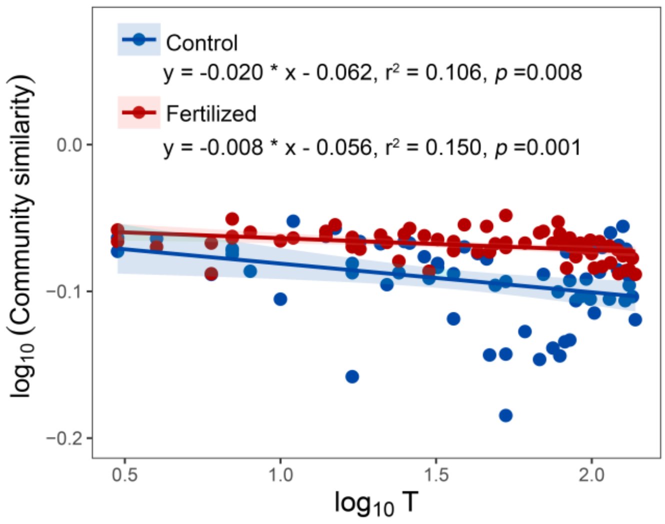


**Fig. S2 Time-decay relationships (TDR) of soil microbial communities**. GeoChip data were normalized according to the basis that gene intensity is proportional to DNA amount used for hybridization. Indices that take account of species abundance between two communities were used.

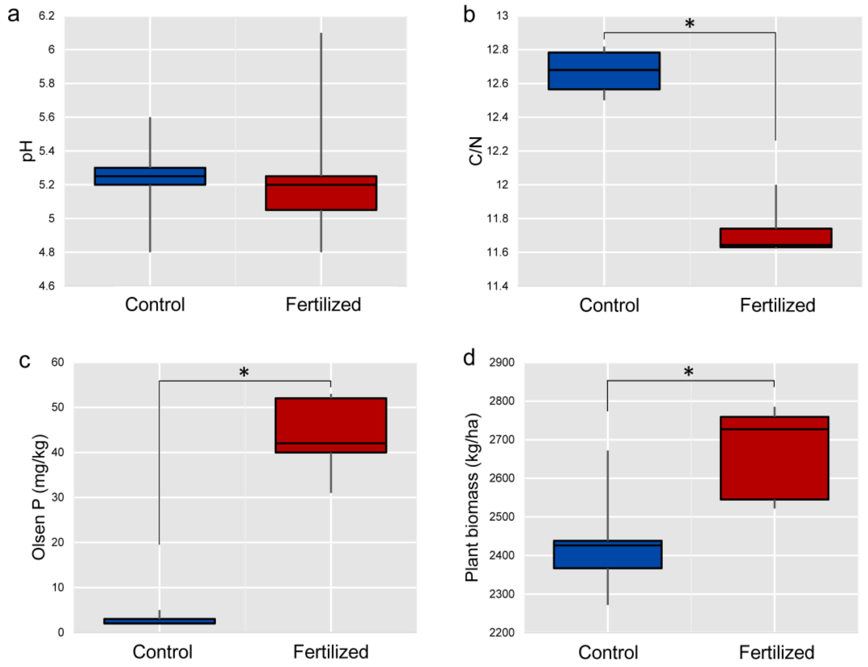


**Fig. S3 Soil geochemical variables and plant biomass in fertilized and control plots.** (a) Soil pH, (b) Soil C:N ratio, (c) Soil Olsen phosphorus concentration, and (d) Plant biomass
